# Opposing roles of ACE2 and TMPRSS2 in IBV Beaudette infection

**DOI:** 10.64898/2026.10.06.753779

**Authors:** Yuan Cao, Yannis Long, Sabrina Schleibinger, Theresa von Heyl, J. J. (Sjaak) de Wit, Benedikt B. Kaufer, Benjamin Schusser, Hicham Sid

## Abstract

Infectious bronchitis virus (IBV) is an economically important avian gammacoronavirus. IBV uses sialic acids and heparan sulfate for cellular attachment, but the host proteins governing its infection remain poorly defined. Here, we investigated the roles of angiotensin-converting enzyme 2 (ACE2) and transmembrane serine protease 2 (TMPRSS2) during IBV Beaudette (IBV BD) infection. ACE2 knockout impaired IBV replication in DF-1 cells, whereas ACE2 reconstitution restored infection. Furthermore, ACE2 overexpression enhanced infection of DF-1 and primary chicken embryo kidney cells, supporting the role of chicken ACE2 as a proviral host factor. In contrast, TMPRSS2 exerted a restrictive function on IBV BD infection as demonstrated in TMPRSS2 KO DF-1 cells and upon pharmacological inhibition, which significantly increased IBV BD infection in DF-1 and Vero cells. RNA sequencing revealed significant changes in lipid metabolism and membrane-related processes following TMPRSS2 knockout and pharmacological inhibition. These findings highlight the opposing roles of ACE2 and TMPRSS2 in IBV BD infection, emphasizing the combined impact of proviral host factors, protease activity, and the cellular membrane environment during gammacoronavirus infections.

## Introduction

Infectious bronchitis virus (IBV) is a member of the genus *Gammacoronavirus* in the order *Nidovirales* and is the primary pathogen responsible for avian infectious bronchitis (Cavanagh, 2007). Highly contagious and pathogenic, IBV poses a devastating threat to the global poultry industry (Jackwood & de Wit, 2020). Since its initial isolation, IBV has diversified into numerous variants characterized by substantial genomic diversity and distinct host-cell preferences, making the comprehensive elucidation of its entry mechanism exceptionally complex (Valastro et al., 2016). Among these variants, Beaudette is a highly attenuated, laboratory-adapted strain whose unusually broad *in vitro* cell tropism was acquired following extensive passage in embryonated chicken eggs. Although it is not representative of circulating field strains, its ability to replicate efficiently in diverse avian and mammalian cell types makes it an ideal experimental model for investigating IBV–host cell interactions (Cunningham et al., 1972; Otsuki et al., 1979).

The specific recognition and binding between the viral spike (S) protein and host-cell receptors are critical determinants of host-cell tropism (Li, 2016; Wickramasinghe et al., 2011). Current research indicates that IBV utilizes specific carbohydrates, such as *α* −2,3-linked sialic acid (SA) and heparan sulfate (HS), as initial attachment receptors; however, tighter binding and subsequent virus-host membrane fusion require the involvement of additional functional core receptors or co-receptors (Madu et al., 2007; Winter et al., 2006). Concurrently, studies have shown that the critical S1/S2 and S2’ cleavage sites within the S proteins of IBV Beaudette strain and SARS-CoV-2 share functional motifs and sequence similarity, both possess a polybasic cleavage site composed of consecutive basic amino acids (e.g., arginine and lysine), as well as a conserved arginine-rich motif at the S2’ site upstream of the fusion peptide (Bestle et al., 2020; Millet & Whittaker, 2015). It has been demonstrated that SARS-CoV-2 utilizes angiotensin-converting enzyme 2 (ACE2) as its primary receptor (Wrapp et al., 2020; Zhou et al., 2020). In the case of IBV, the role of ACE2 remained elusive, while only the upregulation of ACE2 expression in IBV QX-infected tissues has been previously observed (Khan et al., 2025). Therefore, we set out to address the role of ACE2 as a potential receptor during IBV infection of host cells. Furthermore, coronavirus depends not only on S protein-receptor binding but also on S protein activation by host proteases, which primes the spike protein to mediate virus-host cell membrane fusion (Li, 2016). For several mammalian coronaviruses, including SARS-CoV-2, spike-mediated entry can occur at the cell surface. In this pathway, host proteases such as transmembrane serine protease 2 (TMPRSS2) proteolytically prime the spike protein, thereby facilitating fusion of the viral envelope with the plasma membrane (Hoffmann et al., 2020). Alternatively, in the absence of surface proteases, the virus enters the cell via endocytosis, followed by S protein activation mediated by cathepsins within lysosomes or endosomes (Koch et al., 2021). Based on these findings, we hypothesize that IBV may employ a highly sophisticated entry network cooperatively regulated by specific receptors and proteases.

The molecular entry mechanism of this virus, particularly its core host-cell receptors and the activation mechanism of its S protein, remains poorly understood. To fill this knowledge gap, this study aimed to investigate the roles of ACE2 and TMPRSS2 in IBV entry into host cells, thereby providing concrete molecular insights into the host-factor requirements for avian coronavirus infection.

## Results

### Knockout of ACE2 and TMPRSS2 exerts opposing effects on IBV BD infection in DF-1 cells

To investigate the roles of ACE2 and TMPRSS2 during IBV BD infection, we first generated ACE2 and TMPRSS2 knockout (KO) DF-1 cell lines using the CRISPR/Cas9 system (Fig. 1A) (Schleibinger et al., 2026). The ACE2 KO cell line carried a frameshift mutation resulting in a premature stop codon, whereas the TMPRSS2 KO cell line was generated by excising the genomic region encoding the serine protease domain.

**Figure 1.**
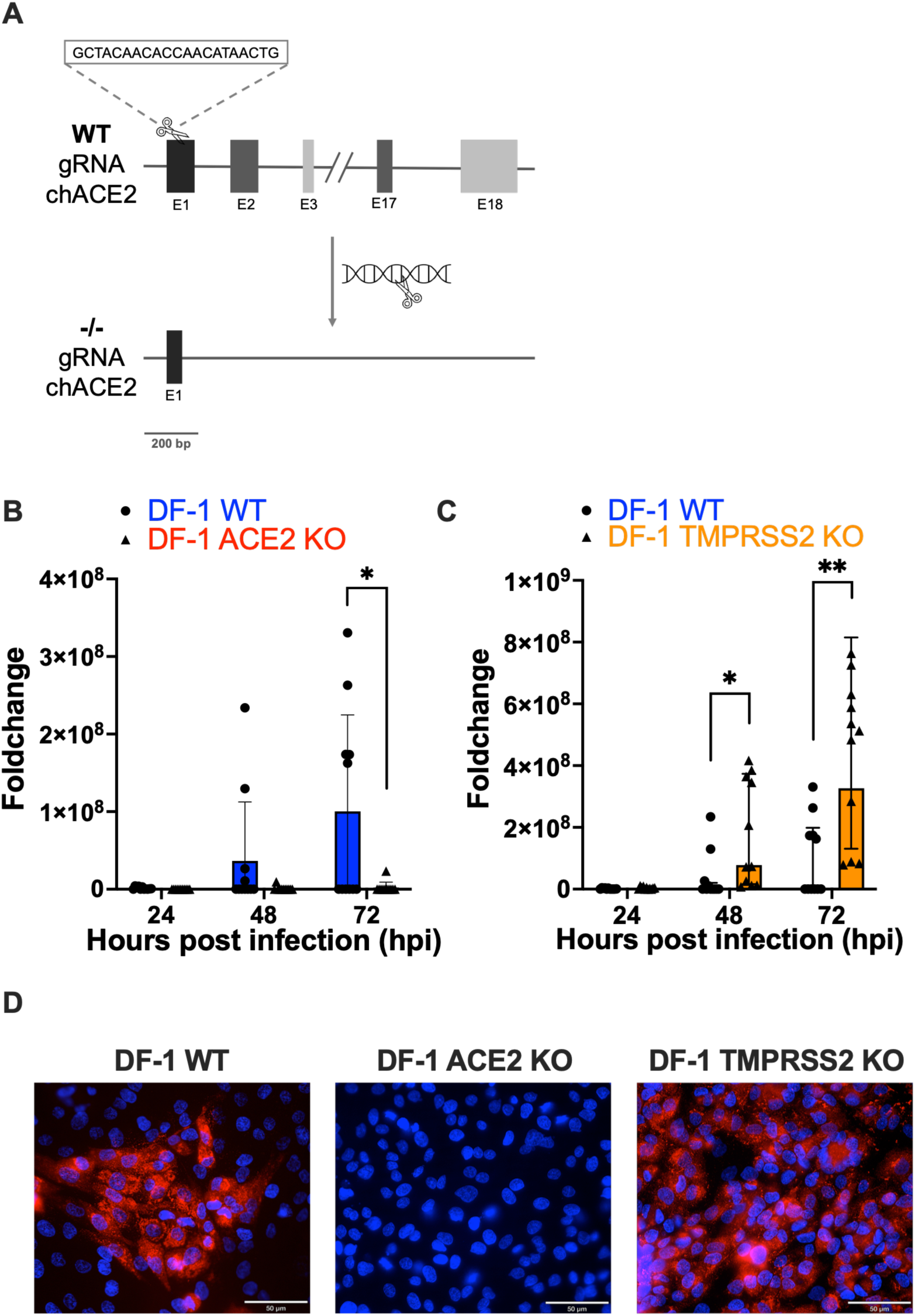
Knockout of ACE2 and TMPRSS2 exerts opposing effects on IBV BD infection in DF-1 cells. (A) Schematic of the CRISPR/Cas9 targeting strategy used to generate ACE2 KO DF-1 cells. The introduced frameshift mutation resulted in a premature stop codon, disrupting the production of functional ACE2 protein. (B) RT-qPCR analysis of IBV BD replication in wild-type (WT) and ACE2 KO cells at 24, 48, and 72 hpi. Viral RNA loads were normalized to 28S expression. All samples were normalized to the WT mock control (*n* = 11). (C) RT-qPCR analysis of IBV BD replication in WT and TMPRSS2 KO cells at 24, 48, and 72 hpi. Viral RNA loads were normalized to 28S expression. All samples were normalized to the WT mock control (*n* = 11). (D) Immunofluorescence assay (IFA) of IBV BD infected WT, ACE2 KO, and TMPRSS2 KO DF-1 cells at 72 hpi. Red signals indicate IBV, and nuclei were counterstained with DAPI (blue). Data are presented as mean ± SD from three independent experiments. Statistical significance was determined using two-sided multiple unpaired t-tests. \**p* < 0.05, \*\**p* < 0.01 compared with the WT control group. Scale bars = 50 μm.

To assess the role of host factors, the generated DF-1 cell lines (WT, ACE2 KO, and TMPRSS2 KO) were infected with IBV BD. Virus replication was subsequently assessed by quantitative reverse transcription PCR (RT-qPCR). At 72 hours post-infection (hpi), IBV BD replication was significantly reduced in ACE2 KO DF-1 cells compared to the WT control group (*p* < 0.05) (Fig. 1B). Conversely, viral replication in TMPRSS2 KO cells was markedly elevated (*p* < 0.01) (Fig. 1C). Consistently, immunofluorescence assays (IFA) at 72 hpi corroborated these findings, revealing a striking decrease in viral positive signals in ACE2 KO cells but a substantial increase in TMPRSS2 KO cells relative to WT controls (Fig. 1D).

Taken together, these data show that ACE2 promotes, whereas TMPRSS2 restricts, IBV BD infection in DF-1 cells.

### Overexpression of chicken ACE2 rescues IBV infection in ACE2 KO cells

To directly confirm that the loss of ACE2 is responsible for the attenuated viral infection and to further evaluate whether ectopic expression of ACE2 can enhance host susceptibility, we performed ACE2 complementation and overexpression experiments. We constructed a plasmid expressing chicken ACE2 with C-terminal HA tag (chACE2-HA) and transfected it into both WT and ACE2 KO DF-1 cells. Subsequent RT-qPCR analysis demonstrated that restoring ACE2 expression in ACE2 KO cells rescued IBV BD infection, resulting in a significant increase in viral loads (*p* < 0.01) (Fig. 2B). Furthermore, ectopic overexpression of chACE2 in WT DF-1 cells also enhanced IBV BD replication (*p* < 0.05) (Fig. 2B). These results were confirmed by IFA, which showed that ACE2 reconstitution restored susceptibility to IBV BD infection (Fig. 2D).

**Figure 2.**
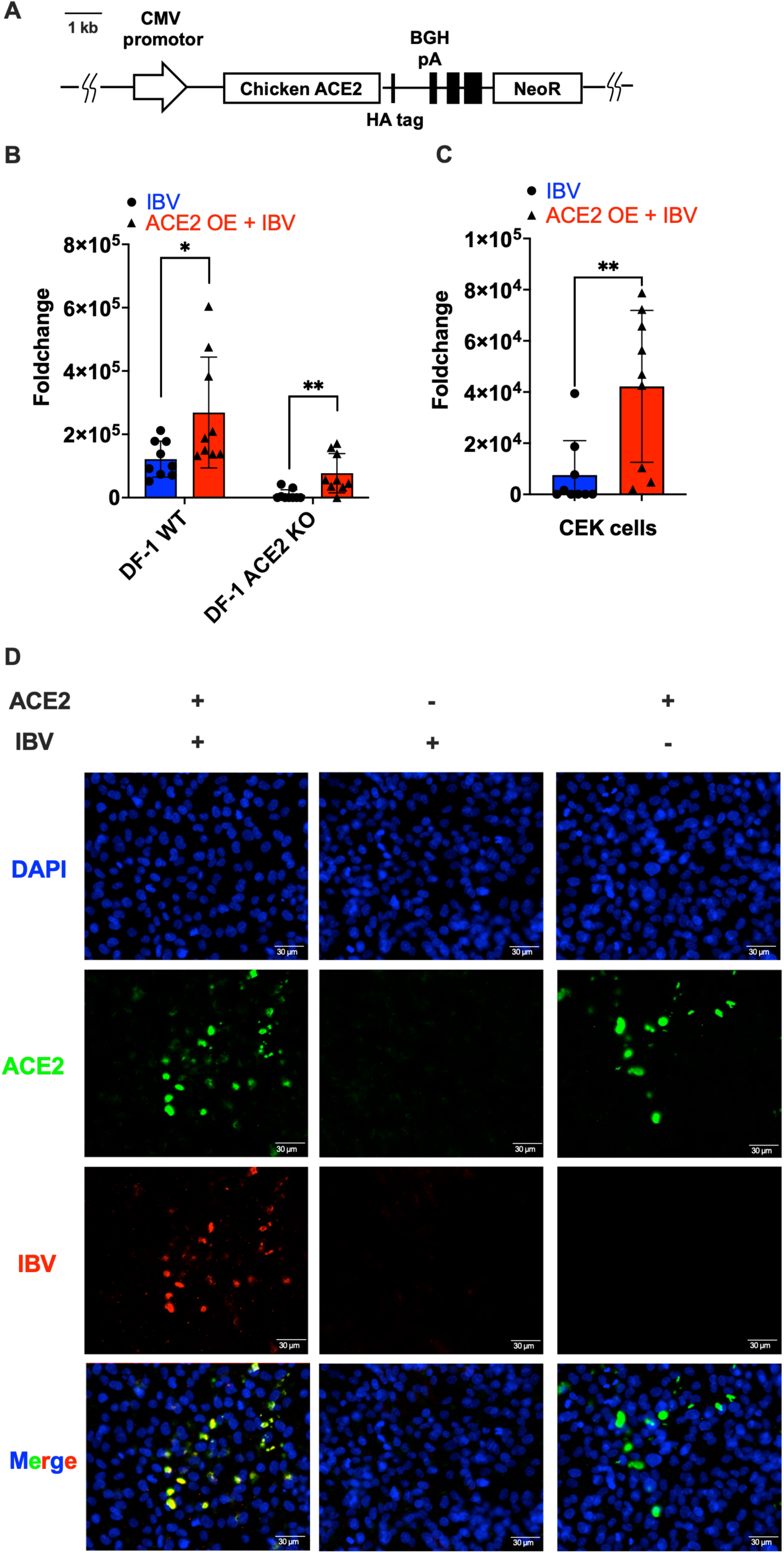
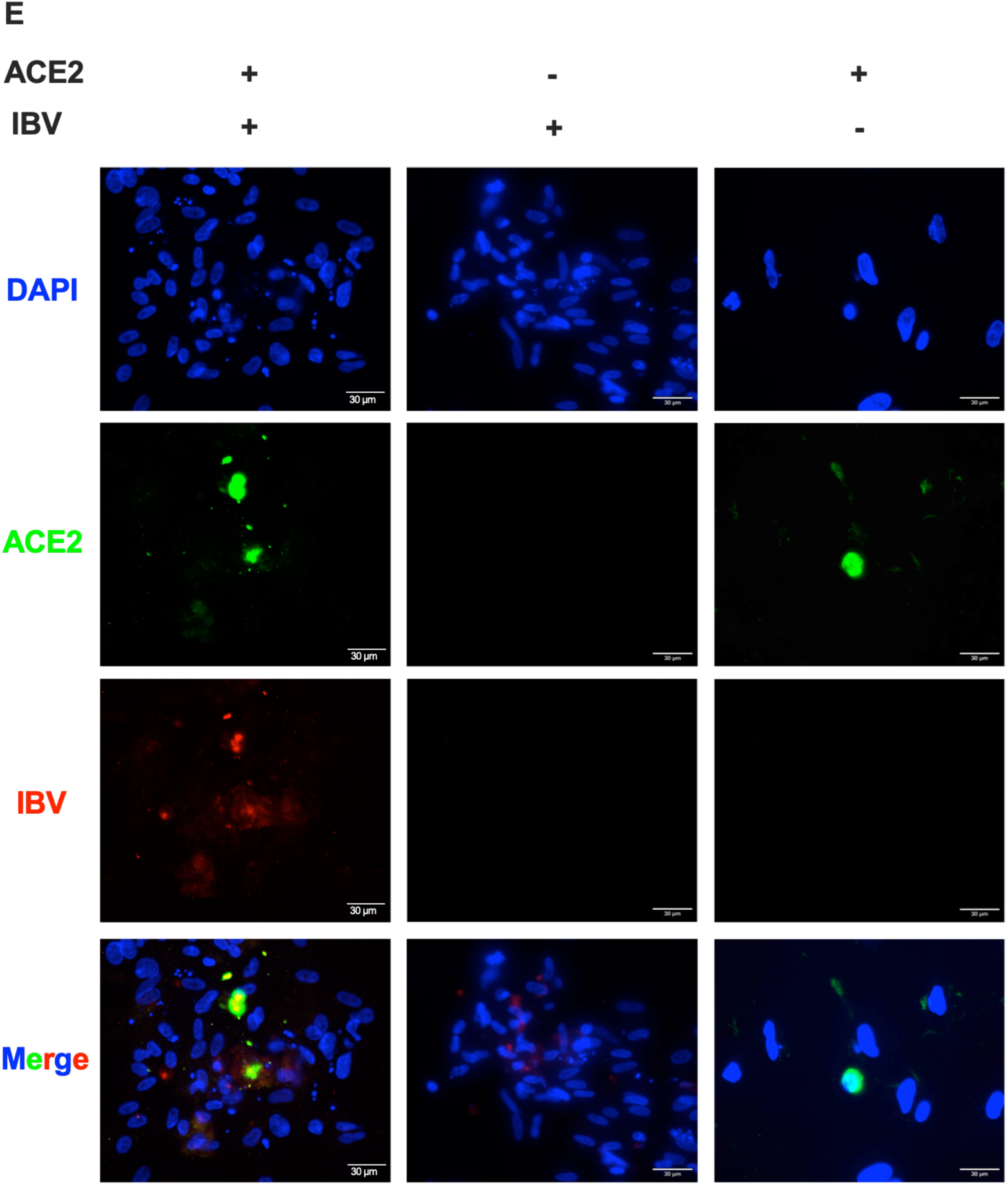
Overexpression of chicken ACE2 rescues and enhances IBV BD infection in avian cells. (A) Schematic diagram of the chACE2 expression plasmid encoding C-terminally HA-tagged chACE2. (B) RT-qPCR analysis of IBV BD replication in WT and ACE2 KO DF-1 cells transfected with chACE2 expression plasmid. All samples were normalized to the WT mock control (*n* = 9). (C) RT-qPCR analysis of IBV BD replication in chicken embryo kidney (CEK) cells transfected with chACE2 expression plasmid. All samples were normalized to the WT mock control (*n* = 9). (D) Representative IFA images showing the restoration and enhancement of IBV BD susceptibility in ACE2 KO DF-1 cells after chACE2 overexpression. (E) Representative IFA images showing the restoration and enhancement of IBV BD susceptibility in CEK cells after chACE2 overexpression. Data are presented as mean ± SD from three independent experiments. Statistical significance was determined using two-sided multiple unpaired t-tests. \**p* < 0.05, \*\**p* < 0.01 compared with the WT control group.

To evaluate the generalizability of these findings in primary avian cells, we overexpressed chACE2 in chicken embryo kidney (CEK) cells. Both RT-qPCR and IFA analyses demonstrated that ACE2 overexpression significantly promoted IBV BD infection in CEK cells (*p* < 0.01) (Fig. 2C, 2E).

We further evaluated the effect of chACE2 overexpression in additional cell models. However, chACE2 overexpression did not significantly enhance IBV BD infection in Vero, HEK293, or Caco-2 cells (Supplementary Fig. S1). These findings indicate that the infection-enhancing effect of chicken ACE2 observed with IBV BD is dependent on the cellular context.

Collectively, these data demonstrate that chicken ACE2 acts as a key functional factor driving IBV BD infection in both immortalized DF-1 cells and primary chicken embryo kidney (CEK) cells.

### Pharmacological inhibition of TMPRSS2 promotes IBV BD replication via a cathepsin-independent endosomal entry pathway

To validate the inhibitory effect of TMPRSS2 on IBV BD infection using pharmacological approaches, DF-1 cells were treated with the TMPRSS2-specific inhibitor Camostat mesylate, and viral replication was monitored by RT-qPCR. Camostat treatment significantly enhanced IBV BD infection at 48 hpi (*p* < 0.01) (Fig. 3A). To determine whether this effect extends to mammalian systems, we validated this approach in Vero cells. Similarly, Camostat treatment markedly accelerated viral replication at 24 hpi (*p* < 0.05) and 72 hpi (*p* < 0.01) (Fig. 3B).

**Figure 3.**
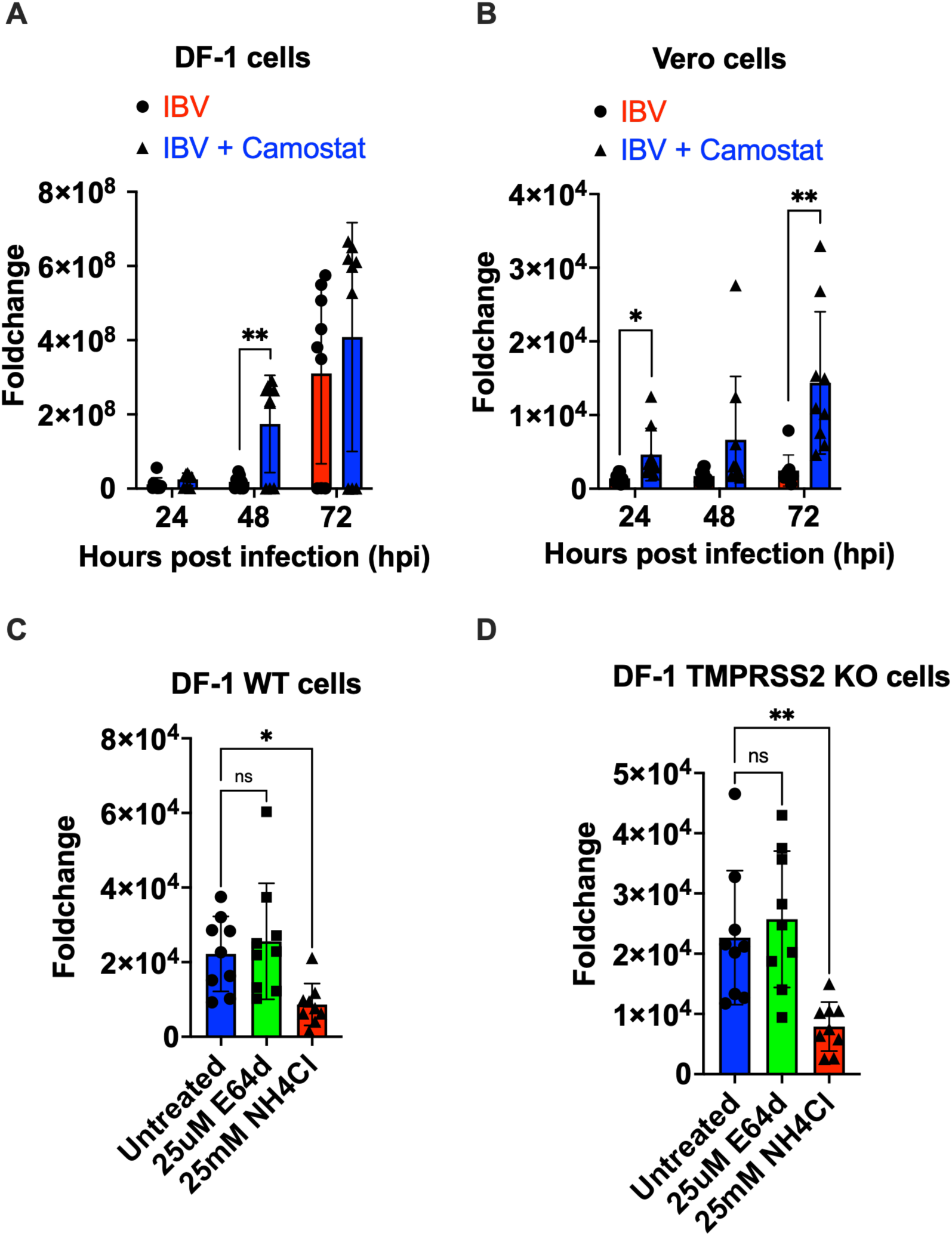
Chemical inhibition of TMPRSS2 increases IBV BD replication via a cathepsin-independent endosomal pathway. (A) RT-qPCR analysis of IBV BD RNA levels in DF-1 cells treated with the TMPRSS2 inhibitor Camostat or DMSO control at 24, 48 and 72 hpi. (B) RT-qPCR analysis of IBV BD replication in mammalian Vero cells treated with Camostat at 24, 48 and 72 hpi. (C, D) Functional analysis of entry pathways. WT and TMPRSS2 KO DF-1 cells were treated with the cathepsin inhibitor E64d or the endosome acidification inhibitor NH_4_Cl prior to IBV BD infection, and viral replication was quantified by RT-qPCR. Data are presented as mean ± SD from three independent experiments. \**p* < 0.05 and \*\**p* < 0.01 compared with the WT control group. Statistical significance was determined using a t-test in A and B, and one-way ANOVA in C and D. All samples were normalized to the WT mock control (*n* = 9).

To determine whether the enhanced IBV BD infection following TMPRSS2 deficiency or inhibition involved an alternative endosomal entry route, we examined the requirements for endosomal acidification and cysteine cathepsin activity. NH_4_Cl was used to neutralize endosomal pH, whereas E64d, a cell-permeable broad-spectrum inhibitor of cysteine cathepsins, including cathepsins B, H and L, was used to assess the requirement for endosomal cysteine-protease activity. Interestingly, E64d treatment failed to suppress IBV BD infection in either WT or TMPRSS2 KO DF-1 cells. In contrast, NH_4_Cl treatment significantly abolished viral infection in both cell lines (*p* < 0.05, *p* < 0.01) (Fig. 3C, 3D), indicating that IBV BD infection requires endosomal acidification but is independent of E64d-sensitive cysteine cathepsin activity under the conditions tested.

Together, these results confirm that pharmacological inhibition of TMPRSS2 promotes IBV BD infection. Furthermore, the differential effects of E64d and NH_4_Cl indicate that IBV BD does not depend on E64d-sensitive cathepsins but requires other acidification-dependent processes within the endosomal pathway.

### TMPRSS2 knockout and pharmacological inhibition induce partially overlapping transcriptional changes in DF-1 cells

To investigate the mechanisms underlying the enhanced IBV infection following TMPRSS2 knockout or pharmacological inhibition, we performed RNA sequencing (RNA-seq) on WT, TMPRSS2 KO, and Camostat-treated DF-1 cells. Each cell group was either uninfected (mock) or infected with IBV BD, resulting in six experimental groups, each with three to four biological replicates. Principal component analysis (PCA) of the batch-corrected expression data showed that biological replicates generally clustered according to experimental group, indicating good within-group reproducibility (Fig. 4A). The first two principal components, PC1 and PC2, which represent the largest and second-largest sources of variation in global gene expression, accounted for 53.6% and 22.4% of the total variance, respectively. TMPRSS2 KO samples separated clearly from both WT and camostat-treated samples along PC1. Thus, TMPRSS2 knockout induced a pronounced global transcriptional shift that was not fully recapitulated by Camostat treatment (Fig. 4A). We therefore focused on the transcriptional changes caused by pharmacological and genetic inhibition of TMPRSS2 to identify candidate molecular processes potentially associated with their shared effects on IBV infection.

**Figure 4.**
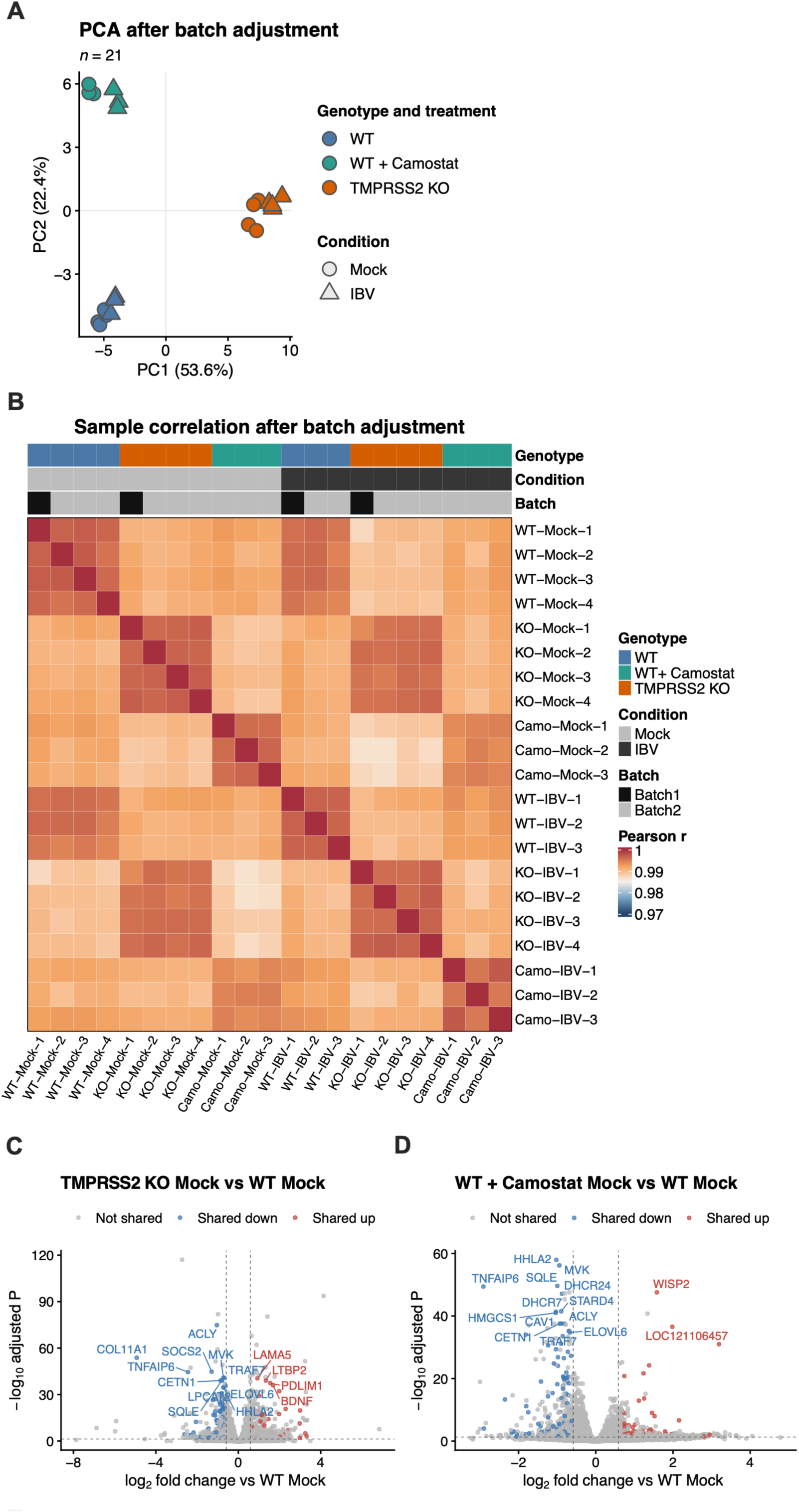
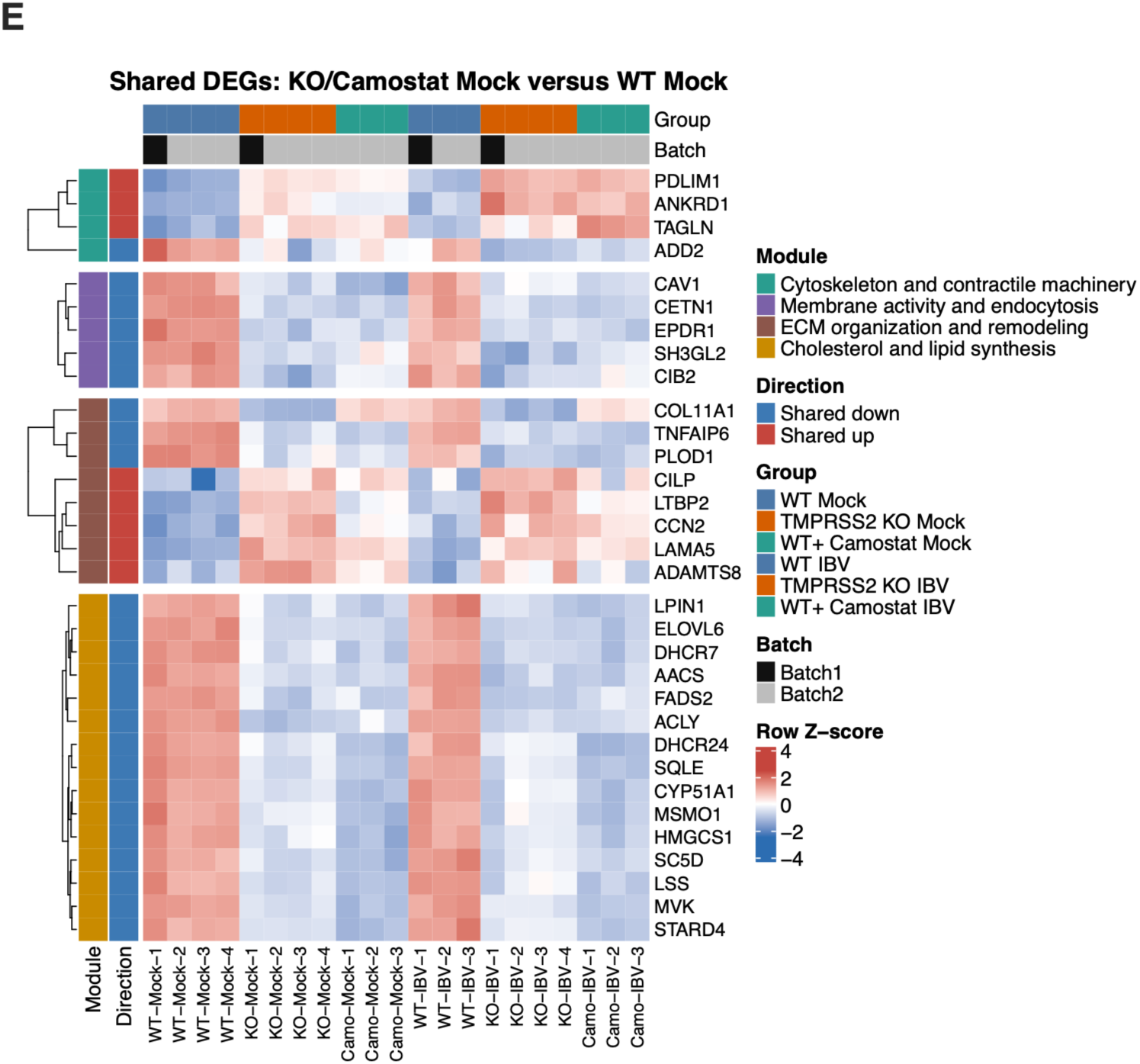
TMPRSS2 knockout and Camostat treatment alter the transcriptomic landscape of DF-1 cells. (A) Principal component analysis (PCA) of RNA-seq samples after batch adjustment. Principal component analysis was performed using the batch-adjusted gene-expression matrix from WT, Camostat-treated, and TMPRSS2 KO DF-1 cells. Each point represents an independent biological sample. (B) Correlation between RNA-seq samples after batch adjustment. Pairwise Pearson correlation coefficients were calculated from the batch-adjusted gene-expression matrix. (C) Volcano plots showing differentially expressed genes in TMPRSS2 KO Mock versus WT Mock cells. (D) Volcano plots showing differentially expressed genes in Camostat Mock versus WT Mock cells. Genes commonly downregulated in both comparisons are shown in blue, genes commonly upregulated are shown in red, and non-shared genes are shown in grey. Differentially expressed genes were defined using an adjusted *P* value of <0.05 and an absolute log₂ fold change of >0.585. Selected representative genes are labeled. (E) Heatmap showing the expression patterns of genes that changed in the same direction in TMPRSS2 KO Mock and Camostat Mock cells relative to WT Mock cells across all six experimental groups. Genes were grouped into functional modules. Expression values are shown as row-scaled Z-scores.

Pearson correlation analysis further showed high overall similarity among the batch-corrected samples, with pairwise correlation coefficients ranging from approximately 0.97 to 1.00 (Fig. 4B). Biological replicates within the same experimental group were generally highly correlated, further supporting the reproducibility of the RNA-seq data. Samples from different batches did not form obvious batch-specific correlation patterns, indicating no prominent residual batch structure in the corrected expression matrix. Differences in correlation remained among genotypes and infection conditions, broadly consistent with the treatment-associated transcriptional variation observed in the PCA.

Differential expression analysis of the uninfected cells identified 493 upregulated and 377 downregulated genes in TMPRSS2 KO Mock cells relative to WT Mock cells. Camostat treatment resulted in 325 upregulated and 353 downregulated genes relative to WT Mock cells, using an adjusted *p*-value <0.05 and an absolute log_2_ fold change >0.585 as the significance thresholds (Fig. 4C, D). Comparison of the two datasets identified 64 commonly upregulated and 95 commonly downregulated genes (Fig. 4C, D).

Selected shared differentially expressed genes were organized into four functional modules: cytoskeletal and contractile regulation, membrane activity and endocytosis, extracellular-matrix organization and remodeling, and cholesterol and lipid metabolism (Fig. 4E). Across all six experimental groups, these genes showed partially similar expression patterns in TMPRSS2 KO Mock and Camostat Mock cells. Thus, although TMPRSS2 KO and Camostat treatment did not produce identical global transcriptomic profiles, they induced partially overlapping basal transcriptional changes involving several cellular programs.

### STRING and GO analyses reveal diverse functional changes following TMPRSS2 knockout

To examine the potential functional relationships among these differentially expressed genes, we constructed a protein association network (Fig. 5A). Sterol-metabolism-associated proteins, including HMGCS1, MVK, SQLE, LSS, MSMO1, CYP51A1, SC5D, DHCR7, and DHCR24, formed a densely connected module. The network also contained extracellular-matrix-associated proteins, including LTBP2, COL11A1, PLOD1, LAMA5, CILP, and ADAMTS8, structural and mechanosensitive proteins, including TAGLN, PDLIM1, and ANKRD1, and membrane-organization or membrane-trafficking-associated proteins, including CAV1 and SH3GL2. These results showed that TMPRSS2-associated perturbations were accompanied by transcriptional changes across multiple cellular programs rather than being restricted to a single metabolic pathway.

**Figure 5.**
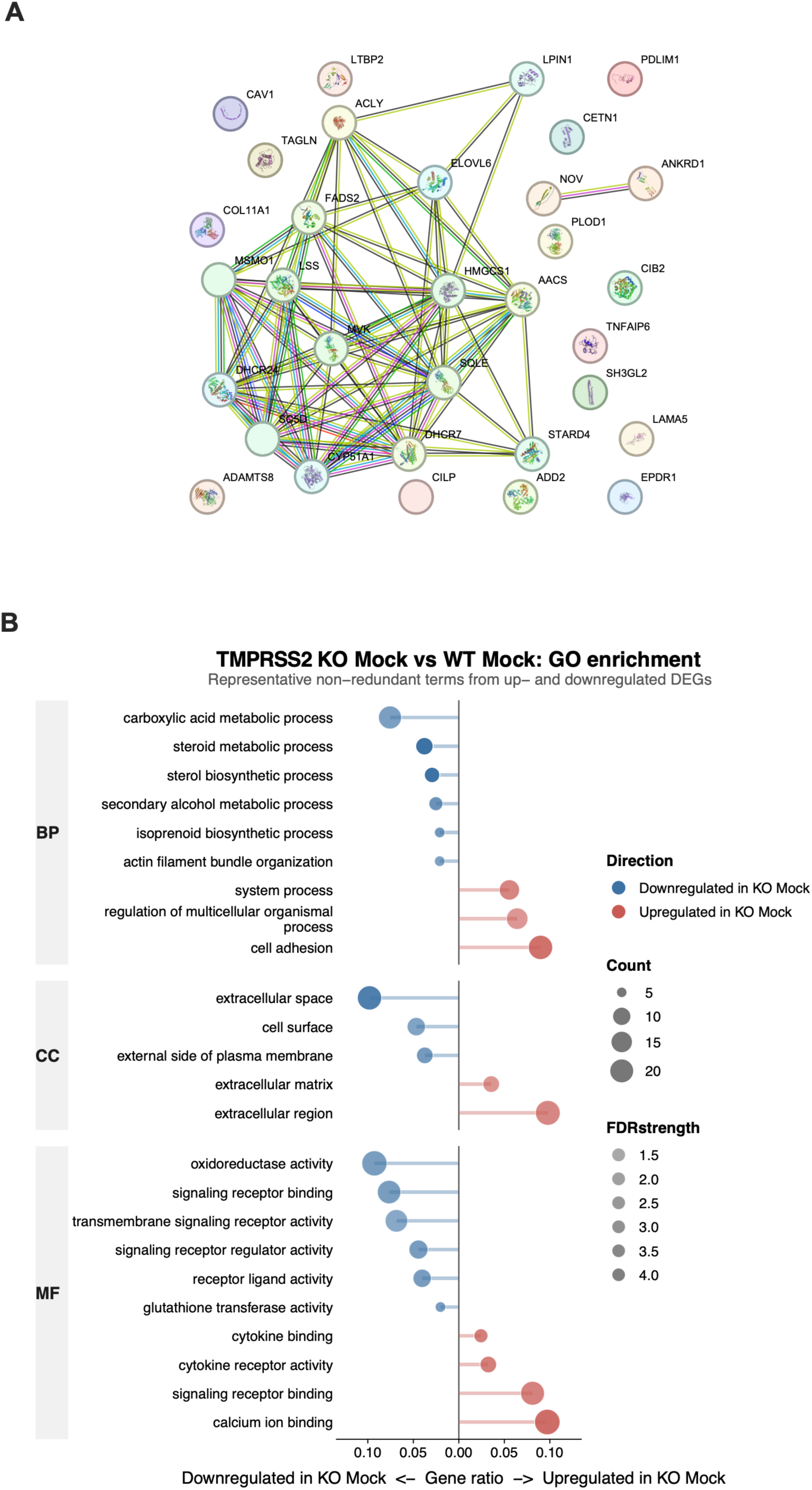
STRING and GO analyses identify multiple cellular programs associated with TMPRSS2 perturbation. (A) STRING protein association network of proteins encoded by selected shared differentially expressed genes. (B) Gene Ontology (GO) enrichment analysis was performed separately for genes upregulated and downregulated in TMPRSS2 KO Mock cells relative to WT Mock cells. Representative non-redundant terms from the biological process (BP), cellular component (CC), and molecular function (MF) categories are shown.

We next performed Gene Ontology (GO) enrichment analysis separately for upregulated and downregulated genes in TMPRSS2 KO Mock cells relative to WT Mock cells (Fig. 5B). This analysis was used to obtain a broader functional overview of the transcriptional consequences of TMPRSS2 knockout. Within the biological process category, downregulated genes were enriched in carboxylic acid metabolism, steroid metabolism, sterol biosynthesis, secondary alcohol metabolism, isoprenoid biosynthesis, and actin filament bundle organization. By contrast, upregulated genes were enriched in cell adhesion, system processes, and the regulation of multicellular organismal processes. Cellular-component analysis associated the downregulated genes with the extracellular space, cell surface, and external side of the plasma membrane, whereas upregulated genes were enriched in the extracellular matrix and extracellular region. Molecular-function analysis further associated the downregulated genes with oxidoreductase activity, transmembrane signaling receptor activity, receptor regulator activity, receptor ligand activity, and glutathione transferase activity. Upregulated genes were enriched for cytokine binding, cytokine receptor activity, signaling receptor binding, and calcium-ion binding (Fig. 5B).

Collectively, the protein association network and GO enrichment analysis identified coordinated changes in lipid and sterol metabolism, extracellular-matrix organization, cell adhesion, cytoskeletal regulation, and receptor-associated signaling. These functional associations were broadly consistent with the expression modules identified in Fig. 4E. These processes represent candidate molecular links between TMPRSS2 perturbation and the enhanced IBV infection phenotype, although their individual contributions require functional validation.

## Discussion

Our study reveals opposing roles for chicken ACE2 and TMPRSS2 during IBV Beaudette infection. ACE2 deficiency markedly reduced viral infection, whereas ACE2 reconstitution or overexpression restored and enhanced viral replication. In contrast, both TMPRSS2 deficiency and pharmacological inhibition promoted viral infection. These findings suggest that the use of host factors by IBV Beaudette differs from the canonical ACE2 receptor–TMPRSS2-mediated fusion paradigm and may reflect distinctive host adaptation strategies of gammacoronaviruses with respect to receptor usage and protease dependence (Hoffmann et al., 2020; Koch et al., 2021).

Previous studies have shown that IBV uses carbohydrate molecules, including α2,3-linked sialic acids and heparan sulfate, to mediate initial attachment (Madu et al., 2007; Winter et al., 2006). However, additional host factors may be required for stable binding, internalization, and membrane fusion. In the present study, ACE2 deficiency reduced viral RNA levels and viral antigen signals in IBV Beaudette-infected DF-1 cells. Re-expression of chicken ACE2 in ACE2-deficient cells restored the infection phenotype, and ACE2 overexpression enhanced IBV infection in both DF-1 and chicken embryo kidney cells. Together, these findings support chicken ACE2 as an important proviral host factor during IBV Beaudette infection. Notably, the current evidence does not clarify whether ACE2 acts as a direct viral attachment receptor, participates in post-attachment entry, or indirectly affects cellular susceptibility by regulating the membrane environment.

In contrast to the proviral role of ACE2, TMPRSS2 acted as a host restriction factor in the cell systems examined. TMPRSS2 KO markedly enhanced IBV Beaudette infection, and pharmacological inhibition with Camostat showed a similar trend in both DF-1 and Vero cells, suggesting that this phenotype is associated with TMPRSS2 activity. This finding contrasts with the well-established role of TMPRSS2 in promoting spike protein activation and cell-surface fusion during infection with SARS-CoV-2 and certain other betacoronaviruses (Hoffmann et al., 2020; Koch et al., 2021; Shirato et al., 2013; Shulla et al., 2011). The divergent effects of TMPRSS2 therefore suggest that its role in coronavirus infection is highly dependent on viral species and cellular context.

Pharmacological experiments further suggested that IBV Beaudette infection depends on endosomal acidification. NH_4_Cl markedly inhibited viral infection in both WT and TMPRSS2-deficient DF-1 cells, whereas E64d showed no substantial inhibitory effect. These results are consistent with previous evidence that IBV can enter host cells through endocytic pathways and suggest that viral infection requires an acidic intracellular environment (Chu et al., 2006; Wang et al., 2019). However, under the experimental conditions used in our study, infection may not depend predominantly on cathepsin activity targeted by E64d.

Transcriptomic analysis of DF-1 cells provided additional clues to the potential cellular basis through which TMPRSS2 restricts IBV Beaudette infection. TMPRSS2 deficiency and Camostat treatment induced highly concordant transcriptional signatures in uninfected DF-1 cells. The most prominent feature was the shared downregulation of genes involved in cholesterol and lipid metabolism, including AACS, ACLY, HMGCS1, MVK, SQLE, LSS, CYP51A1, MSMO1, SC5D, DHCR24, DHCR7, and STARD4, together with genes associated with fatty acid elongation, fatty acid desaturation, and glycerolipid metabolism, including ELOVL6, FADS2, and LIPIN1 (Csaki & Reue, 2010; Guillou et al., 2010; Luo et al., 2020). These findings suggest that TMPRSS2 deficiency or pharmacological inhibition remodels basal lipid-homeostasis programs, potentially altering membrane properties that regulate viral uptake at the plasma membrane and subsequent trafficking or fusion within acidified endosomes.

The coordinated downregulation of these genes spans multiple steps in mevalonate production, sterol biosynthesis, terminal cholesterol modification, intracellular cholesterol transport, fatty acid metabolism, and phospholipid metabolism, suggesting suppression of an SREBP2-associated sterol-biosynthetic transcriptional program (Luo et al., 2020). However, reduced expression of cholesterol-biosynthetic genes should not be interpreted simply as reduced total cellular cholesterol. SREBP2 activation is negatively regulated by free cholesterol and oxysterol levels in the endoplasmic reticulum. When cells sense sufficient sterol levels, the SCAP–INSIG complex limits SREBP2 activation, thereby suppressing the expression of multiple sterol-biosynthetic genes (Brown & Goldstein, 1997; Luo et al., 2020; Radhakrishnan et al., 2008). The observed transcriptional signature may therefore reflect redistribution among intracellular sterol pools, oxysterol accumulation, altered cholesterol transport, or adaptation to exogenous lipid use in the culture system, rather than global cholesterol depletion.

This distinction is particularly important for interpreting the enhanced IBV infection phenotype. IBV Beaudette entry depends on lipid-raft integrity and clathrin-mediated endocytosis (Wang et al., 2019). In this study, acute extraction of plasma-membrane cholesterol with methyl-β-cyclodextrin (MβCD) reduced viral entry. However, acute depletion of plasma-membrane cholesterol by MβCD is not equivalent to the steady-state adaptation induced by long-term TMPRSS2 deficiency or pharmacological inhibition, as cyclodextrins can perturb cholesterol distribution across membrane compartments and may also affect other membrane lipids (Zidovetzki & Levitan, 2007). Cholesterol in the endoplasmic reticulum, plasma membrane, endosomes, lysosomes, and lipid droplets may change independently (Luo et al., 2020; Maxfield & van Meer, 2010). Consequently, downregulation of the cholesterol-biosynthetic transcriptional program does not exclude remodeling of accessible cholesterol, lipid composition, or membrane fluidity within membrane compartments relevant to viral entry.

In addition to altered lipid homeostasis, the coordinated downregulation of genes associated with vesicular activity and endocytosis suggests remodeling of membrane-trafficking networks. CAV1, CETN1, EPDR1, SH3GL2 and CIB2 were reduced in both TMPRSS2-deficient and Camostat-treated cells. CAV1 is a major component of caveolae and cholesterol-rich lipid rafts, whereas SH3GL2 encodes endophilin A1, which participates in membrane curvature and vesicle formation (Boucrot et al., 2015; Parton & Simons, 2007). EPDR1 is associated with lysosomal function and cell adhesion, while CETN1 and CIB2 suggest potential changes in Ca²⁺-dependent membrane dynamics and cellular structural regulation (Dal Cortivo et al., 2019; Wei et al., 2019). Given that IBV Beaudette primarily enters cells via clathrin-mediated endocytosis, these changes may reflect a rebalancing among distinct endocytic pathways, membrane microdomains, and endosomal-trafficking processes, rather than a general increase in endocytic activity (Wang et al., 2019).

Furthermore, the shared upregulation of extracellular matrix (ECM) organization and remodeling genes, including CILP, LTBP2, CCN2, LAMA5 and ADAMTS8, suggests alterations in ECM composition, matrix remodeling and cellular adhesion. The ECM–integrin axis not only determines cell morphology and mechanical properties but can also regulate the spatial distribution of plasma-membrane receptors, membrane-microdomain stability and endocytic activity (Echarri et al., 2007; Kechagia et al., 2019). ECM remodeling may therefore indirectly influence IBV attachment or subsequent entry by altering the local microenvironment at the cell surface.

The increased expression of PDLIM1, ANKRD1 and TAGLN further indicates remodeling of the cytoskeleton and contractile machinery. Actin networks contribute to the maintenance of plasma-membrane tension, the formation of endocytic vesicles, and endosomal trafficking (Kaksonen & Roux, 2018; Lappalainen et al., 2022; Soldati & Schliwa, 2006). These changes may therefore act in concert with altered membrane lipids and vesicular-trafficking networks.

Overall, the RNA-seq data support a model in which TMPRSS2 deficiency enhances IBV Beaudette infection by establishing a membrane-remodeled cellular state that favors productive endosomal infection. TMPRSS2 knockout and Camostat treatment shared transcriptional changes in genes involved in sterol and lipid homeostasis, membrane organization, vesicular trafficking, extracellular-matrix remodeling, and cytoskeletal regulation. In particular, the coordinated downregulation of sterol-biosynthetic genes, together with changes in CAV1, STARD4, SH3GL2, CIB2 and EPDR1, suggests altered organization and intracellular distribution of membrane lipids rather than simply a reduction in total cellular cholesterol. Concurrent changes in extracellular-matrix and cytoskeletal genes may further modify plasma-membrane tension, the spatial organization of attachment factors and the formation or trafficking of endocytic vesicles. We therefore propose that TMPRSS2 deficiency increases the efficiency with which IBV particles are internalized and delivered to acidified endosomal compartments that support membrane fusion. This model is consistent with the strong inhibition produced by NH_4_Cl and the absence of inhibition by E64d, suggesting that the enhanced infection is associated with membrane and endosomal remodeling rather than increased activity of E64d-sensitive cysteine cathepsins. Thus, TMPRSS2 appears to restrict IBV Beaudette infection indirectly by maintaining membrane lipid homeostasis and trafficking, whereas its loss creates a cellular environment that is more permissive to acidification-dependent endosomal infection.

Taken together, genetic knockout, rescue experiments, overexpression, and pharmacological evidence establish a framework in which ACE2 and TMPRSS2 exert opposing functions during IBV Beaudette infection. These findings further suggest that receptor usage, protease activity, and membrane homeostasis may collectively shape cellular susceptibility and entry pathways of gammacoronaviruses.

## Materials and Methods

### Key resources table

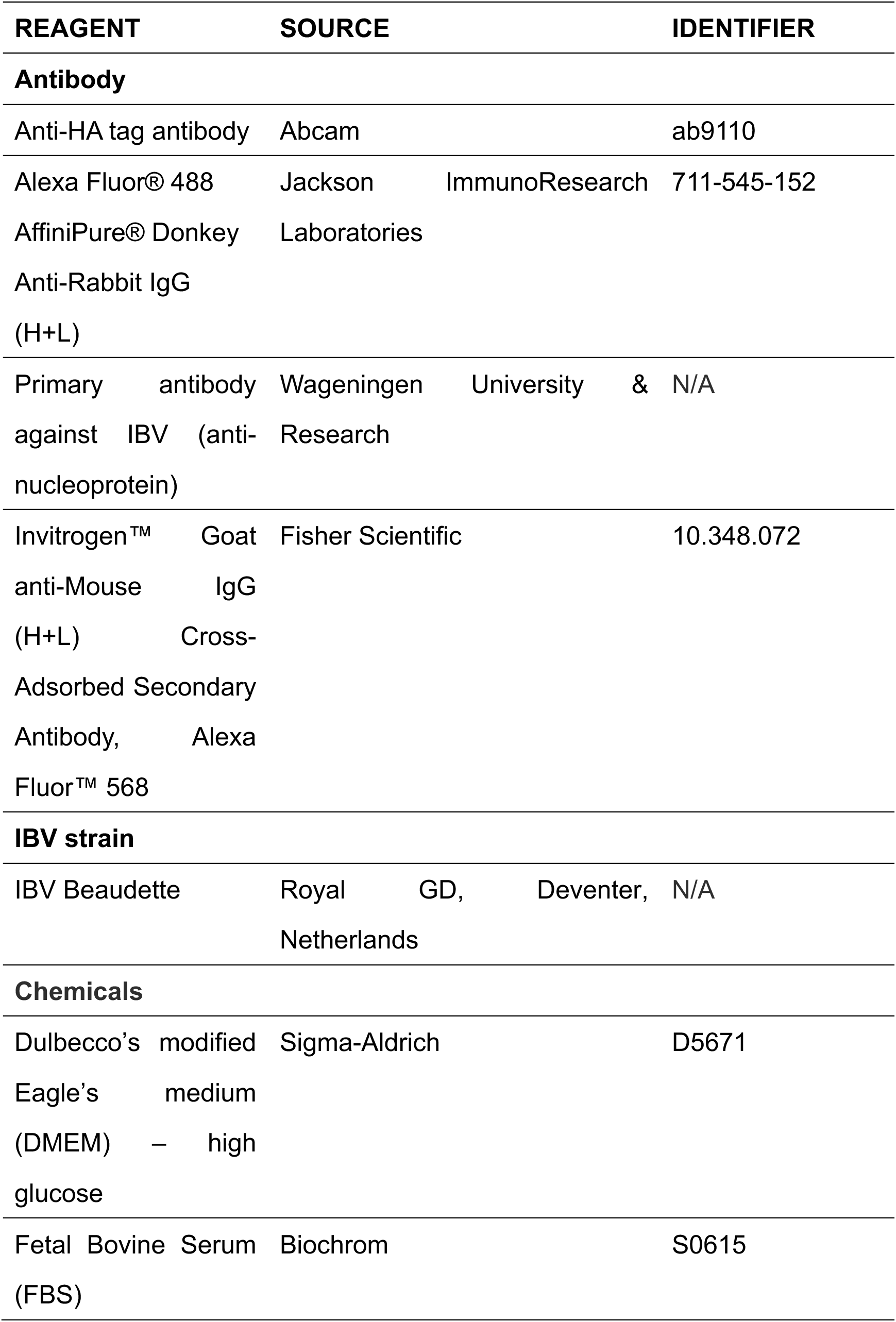
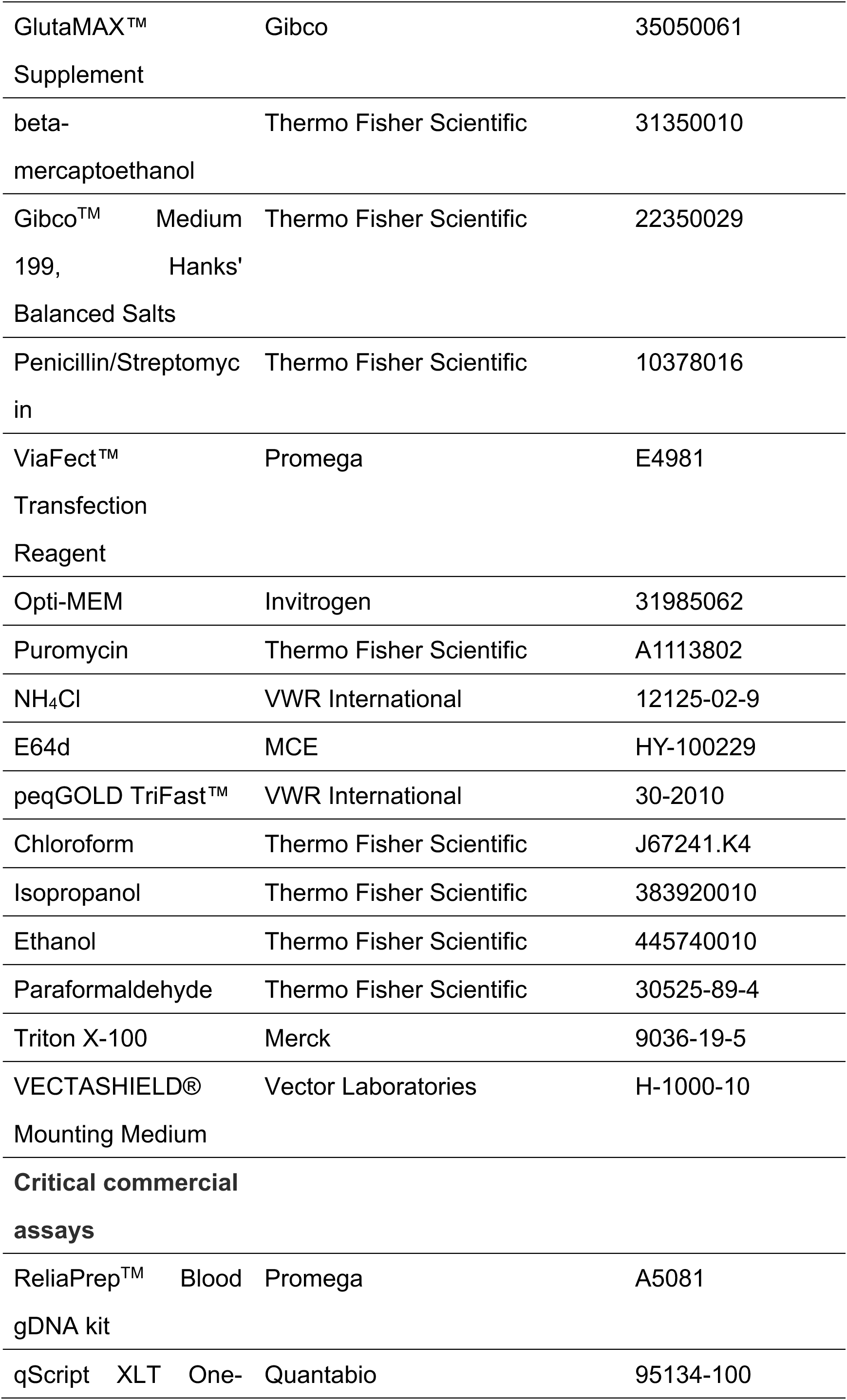
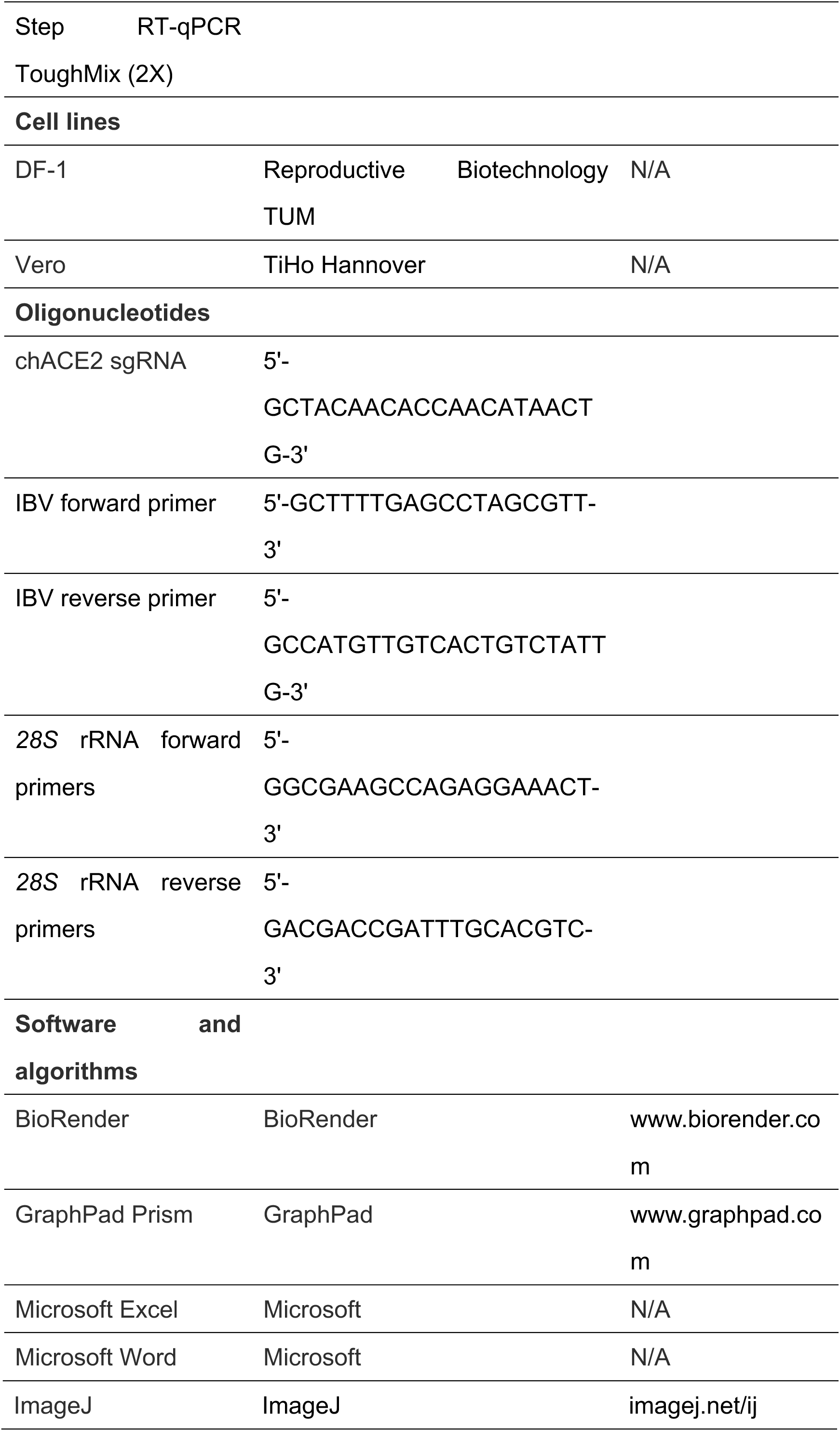

### Cell line

DF-1 (UMNSAH/DF-1) cells cell line, a continuous chicken embryo fibroblast cell line, was obtained from the Friedrich-Loeffler-Institut, Riems, Germany, and is commercially available (CRL-3586, ATCC, VA, USA). They were grown in Dulbecco’s modified Eagle’s medium (DMEM) high glucose (SIGMA, cat # D5671) supplemented with 10% fetal bovine serum (FBS) (Biochrom, cat # S0615) and 1% GlutaMAX™ Supplement (Gibco, cat # 35050061) at 40 °C with 5% CO_2_. The continuous mammalian Vero cell line (derived from the kidney tissue of an adult African green monkey) was kindly provided by Prof. Dr. Silke Rautenschlein, University of Veterinary Medicine, Hannover, and grown in DMEM high glucose supplemented with 10% FBS, 1% GlutaMAX™ Supplement, and 0.2% beta-mercaptoethanol (Thermo Fisher Scientific, cat # 31350010) at 37 °C with 5% CO_2_.

### Isolation of primary chicken embryo kidney cells

Chicken embryos were incubated for 19 days at 37 °C in a ProCon incubator. The embryos were immediately euthanized and the kidneys were carefully isolated, placed into a conical flask, and washed with PBS via gentle agitation on a magnetic stirrer for 5 min. The use of embryonated eggs prior to hatch was in accordance with the German Animal Welfare Act. The permission for animal experiment was not required as indicated by the Animal Welfare Act §8 and §14 of the Animal Welfare Laboratory Animal Regulation (Tierschutz-Versuchstierverordnung (TierSchVersV)). For tissue dissociation, the kidneys were subjected to trypsinization at 37 °C with continuous magnetic stirring. After 30 min, the digestion was quenched by adding 50 mL of complete culture medium to the turbid cell suspension. The suspension was filtered through a cell strainer, collected into 50 mL centrifuge tubes, and centrifuged at 300 x *g* for 10 min. The supernatant was discarded, and the resulting cell pellet was passed through a cell strainer once more to obtain a single-cell suspension. Finally, the cells were resuspended and cultured in complete growth medium consisting of 500 mL Gibco Medium 199 (with Hank’s Salts, L-Glutamine, 25 mM HEPES, and L-Amino Acids) (Thermo Fisher Scientific, cat # 22350029), supplemented with 25 mL FBS, 5 mL GlutaMAX™, and 5 mL Penicillin/Streptomycin (Thermo Fisher Scientific, cat # 10378016).

### CRISPR/Cas9-mediated knockout of *ACE2*

ACE2 KO DF-1 cells were generated using the pSpCas9(BB)-2A-Puro (PX459) V2.0 plasmid. This plasmid was a gift from Feng Zhang (Addgene plasmid no. 62988; http://n2t.net/addgene:62988; RRID: Addgene_62988). An sgRNA targeting exon 1 of chicken *ACE2* (5′-GCTACAACACCAACATAACTG-3′) was cloned into the PX459 plasmid. Transfection of cells was performed using ViaFect™ Transfection Reagent (Promega, cat # E4981). Briefly, 500 ng of the DNA plasmids and 3 µL ViaFect™ Transfection Reagent were mixed in 50 µL Opti-MEM (Invitrogen, cat # 31985062) and applied to DF-1 cells. The medium was exchanged after 24 hours of incubation, and the transfected cells were selected with puromycin (2.5 μg/mL) (Thermo Fisher Scientific, cat # A1113802). After 48 h the puromycin selection was completed, and the medium was exchanged. The initial pool of transfected cells was expanded and tested via sequencing for genetic modification, and if successful, a limiting dilution was performed to generate a single-cell clone of transfected cells. Genomic DNA was extracted from monoclonal cells using Promega Relia Prep Blood gDNA kit (Promega, cat # A5081) and sequenced to verify the modification of the *ACE2* gene. The final selected positive clone harbored a frameshift mutation in the first exon of *ACE2*, resulting in a premature stop codon. This mutation caused a shift in the open reading frame (ORF), leading to early termination of functional protein translation.

### Virus propagation and titration in chicken embryos

IBV Beaudette was propagated in 10-day-old embryonated chicken eggs via the allantoic route (100 µL/egg). Allantoic fluid was harvested at 3 days post-infection (dpi) and then centrifuged to remove debris. Viral titers were determined by the 50% embryo infectious dose assay, in which specific embryo mortality observed after 24 h was recorded and analyzed (Kint et al., 2015).

### Infection of cells

The cells were seeded 24 h prior to infection. The IBV Beaudette virus stock was thawed on ice and diluted in the culture medium. The cells were washed with PBS before infection. For the standard infection, cells were inoculated at a multiplicity of infection (MOI) of 0.01 for 2 h to allow virus adsorption. The inoculum was then removed, the cells were washed with PBS, and fresh maintenance medium was added until cell collection. Alternatively, for the investigation of the effect of the endosomal system on virus infection, cells were pre-incubated with culture medium containing NH_4_Cl (VWR International, cat # 12125-02-9) or E64d (MCE, cat # HY-100229) for 1 h, followed by infection with the virus at an MOI of 0.1 for 1 h. After this initial infection period, the medium was replaced with fresh drug-containing maintenance medium for an additional 1 h of incubation, then switched to drug-free maintenance medium, and cells were harvested 8 hpi.

### Plasmid transfection and rescue assay

To assess the role of chicken ACE2 during IBV infection, DF-1 and CEK cells were transfected with the ACE2-HA expression plasmid using ViaFect™ Transfection Reagent (Promega, cat # E4981). Briefly, 500 ng of the DNA plasmids and 3 µL ViaFect™ Transfection Reagent were mixed in 50 µL Opti-MEM (Invitrogen, cat # 31985062) applied to DF-1 cells, in a ratio of 6:1. The mixture was incubated at room temperature for 20 min. Before transfection, the cell culture medium was replaced with fresh medium, and the transfection complexes were added to the cells. Viral infection was performed 48 h post-transfection.

### RNA isolation

Total RNA was isolated using peqGOLD TriFast™ (VWR International, cat # 30-2010) according to the manufacturer’s instructions with minor modifications. Briefly, culture medium was removed and 500 µL peqGOLD TriFast™ was added to lyse the cells in 6-well plates. The lysate was mixed thoroughly with 100 µL of chloroform (Thermo Fisher Scientific, cat # J67241.K4) and incubated at room temperature for 5 min. Following centrifugation at 12,000 × *g* for 6 min, the upper aqueous phase was transferred to a fresh tube and mixed with 250 µL isopropanol (Thermo Fisher Scientific, cat # 383920010). The mixture was incubated on ice for 10 min and centrifuged at 12,000 × *g* for 10 min at 4 °C. The RNA pellet was washed three times with 1 mL 75% ice-cold ethanol (Thermo Fisher Scientific, cat # 445740010). Finally, the pellet was air-dried at room temperature for 60 min and resuspended in 100 µL of nuclease-free water.

### RT-qPCR quantification of viral load

Before the reaction, RNA samples were normalized to a concentration of 20 ng/µL. The 10 µL reaction mixture consisted of 5 µL of qScript XLT One-Step RT-qPCR ToughMix (2X) (Quantabio, cat # 95134-100), 0.5 µL 10 pmol IBV forward and reverse primers (5’-GCTTTTGAGCCTAGCGTT-3’ and 5’-GCCATGTTGTCACTGTCTATTG-3’), 0.5 µL 10 pmol *28S* rRNA forward and reverse primers (5’-GGCGAAGCCAGAGGAAACT-3’ and 5’-GACGACCGATTTGCACGTC-3’), 0.125 µL each of the probe (5 pmol), 1.75 µL of nuclease-free water, and 1 µL of RNA template. For normalization, *28S* rRNA was used as the housekeeping gene for DF-1 cells.

Thermal cycling was performed as follows: reverse transcription at 50 °C for 10 min; initial denaturation at 95 °C for 1 min; followed by 40 cycles of denaturation at 95 °C for 1 min and annealing/extension at 60 °C for 1 min.

Relative viral copy numbers were calculated using the 2^-ΔΔCt^ method. For mock-infected controls, where no viral amplification occurred, a theoretical Ct value of 40 (the maximum cycle number) was assigned. The ΔCt for the mock control was determined by subtracting the average housekeeping gene Ct of all samples from the theoretical value of 40.

### Immunofluorescence staining

Cells were seeded on coverslips in 12-well plates 24 h prior to infection. At the designated time points post-infection, cells were fixed with ice-cold 4% paraformaldehyde (PFA) (Thermo Fisher Scientific, cat # 30525-89-4) for 5 min at room temperature. Following washing with PBS, cells were blocked and incubated with Anti-HA tag antibody (detecting ACE2) (10 μg/mL) (Abcam, cat # ab9110) for 1 h at room temperature in the dark to detect surface protein expression. Cells were washed with PBS and incubated with Alexa Fluor® 488 AffiniPure® Donkey Anti-Rabbit IgG (H+L) (1.7 μg/mL) (Jackson, cat # 711-545-152) for 1 h at room temperature. After the surface staining, cells were permeabilized with 0.1% Triton X-100 in PBS for 5 min. Cells were then washed and incubated with the primary antibody against IBV (anti-nucleoprotein) (20 μg/mL) (Wageningen University & Research) followed by the Invitrogen™ Goat anti-Mouse IgG (H+L) Cross-Adsorbed Secondary Antibody, Alexa Fluor™ 568 (10 μg/mL) (Fisher Scientific, cat # 10.348.072). Coverslips were washed three times with PBS between each antibody incubation step. Finally, coverslips were mounted on glass slides using VECTASHIELD® Mounting Medium (Vector Laboratories, cat # H-1000-10). The slides were subsequently visualized under a fluorescence microscope, Echo Revolve (Discover Echo).

### RNA sequencing and bioinformatic analysis

WT, TMPRSS2 KO, and Camostat-treated DF-1 cells were mock-treated or infected with IBV at an MOI of 0.01. Total RNA was collected at 48 hours post-infection using peqGOLD TriFast™ (VWR International, cat # 30-2010). A total of 21 biologically independent samples were analyzed: WT Mock (*n* = 4), TMPRSS2 KO Mock (*n* = 4), Camostat Mock (*n* = 3), WT IBV (*n* = 3), TMPRSS2 KO IBV (*n* = 4), and Camostat IBV (*n* = 3). RNA quality was assessed using Agilent 2100 Bioanalyzer, and all samples had RNA integrity numbers (RINs) greater than 9. RNA was submitted for Plasmidsaurus RNA-seq (3′ end counting, Illumina NovaSeq X Plus). FASTQ files were processed using the RNA-seq alignment -STAR Pipeline 2.7.10a in the Cancer Genomics cloud platform. RNA-seq reads were aligned to the *Gallus gallus* (chicken) GRCg7b reference genome (Accession Number GCF_016699485.2). The pipeline generates molecular reads per gene, metrics and an alignment file.

All downstream statistical analyses and visualizations were performed in R version 4.6.1. Principal component analysis and Pearson correlation analysis were performed using batch-adjusted VST expression values. Differential expression analysis was conducted using DESeq2 version 1.52.0. The statistical model included batch, genotype, treatment, and the genotype-by-treatment interaction term. TMPRSS2 KO Mock and Camostat Mock samples were independently compared with WT Mock samples. Genes with a Benjamini–Hochberg-adjusted P value of <0.05 and an absolute log₂ fold change of >0.585 were considered differentially expressed. Genes significantly altered in the same direction in both comparisons were defined as shared differentially expressed genes.

Expression patterns of the shared differentially expressed genes across all six experimental groups were visualized using row-scaled Z-scores based on batch-adjusted variance-stabilized expression values. Gene Ontology enrichment analysis was performed separately for genes upregulated and downregulated in TMPRSS2 KO Mock cells relative to WT Mock cells using the R package clusterProfiler and the chicken annotation database org.Gg.eg.db. All DESeq2-tested genes with non-missing adjusted P values and successful mapping to chicken Entrez Gene IDs were used as the background set. GO terms with a Benjamini–Hochberg-adjusted FDR < 0.05 were considered significant. Redundant GO terms were reduced using the simplify function in clusterProfiler with the Wang semantic similarity measure and a similarity cutoff of 0.70. Biological Process, Cellular Component, and Molecular Function categories were analyzed separately and integrated for visualization. Protein functional associations were analyzed using the STRING database, with *Gallus gallus* specified as the organism.

### Quantification and statistical analysis

Statistical analysis was performed using two-sided multiple unpaired *t*-tests for comparisons between two groups and one-way analysis of variance (ANOVA) followed by Dunnett’s multiple-comparisons test for comparisons among more than two groups. *P* values less than 0.05 were considered significant, with *P* < 0.05 and *P* < 0.01 indicated as ^∗^ and ^∗∗^ respectively, ns indicated no significant differences.

## Acknowledgements

The authors thank all the members of the Chair of Reproductive Biotechnology at the TUM School of Life Sciences for their technical assistance and helpful discussions. ChatGPT (https://chat.openai.com/) was used to improve conciseness of language and coding, with no contribution to the scientific content.

## Funding

The project was financed by a grant from the German Research Foundation to HS (Project number: 572662026).

## Author contributions

YC, YL, and SSc: Data curation, Formal analysis, Investigation, Methodology, Writing – review & editing. YC: Writing – original draft. TvH: Data curation, Writing – review & editing. JJW: Conceptualization, Writing – review & editing. BBK: Conceptualization, Writing – review & editing. BS: Conceptualization, Writing – review & editing. HS: Conceptualization, Supervision, Funding acquisition, Writing – review & editing. All authors read, critically reviewed, and approved the final version of the manuscript for submission.

## Conflict of interest

The authors declare that they have no competing interests.

## Data availability

Data supporting the findings of this study are included in the article and its Supplementary Information.

## Notes

### Competing Interest Statement

The authors have declared no competing interest.

