## Supplementary Fig. S1 for "Opposing roles of ACE2 and TMPRSS2 in IBV Beaudette infection"

**This file includes Figures S1**

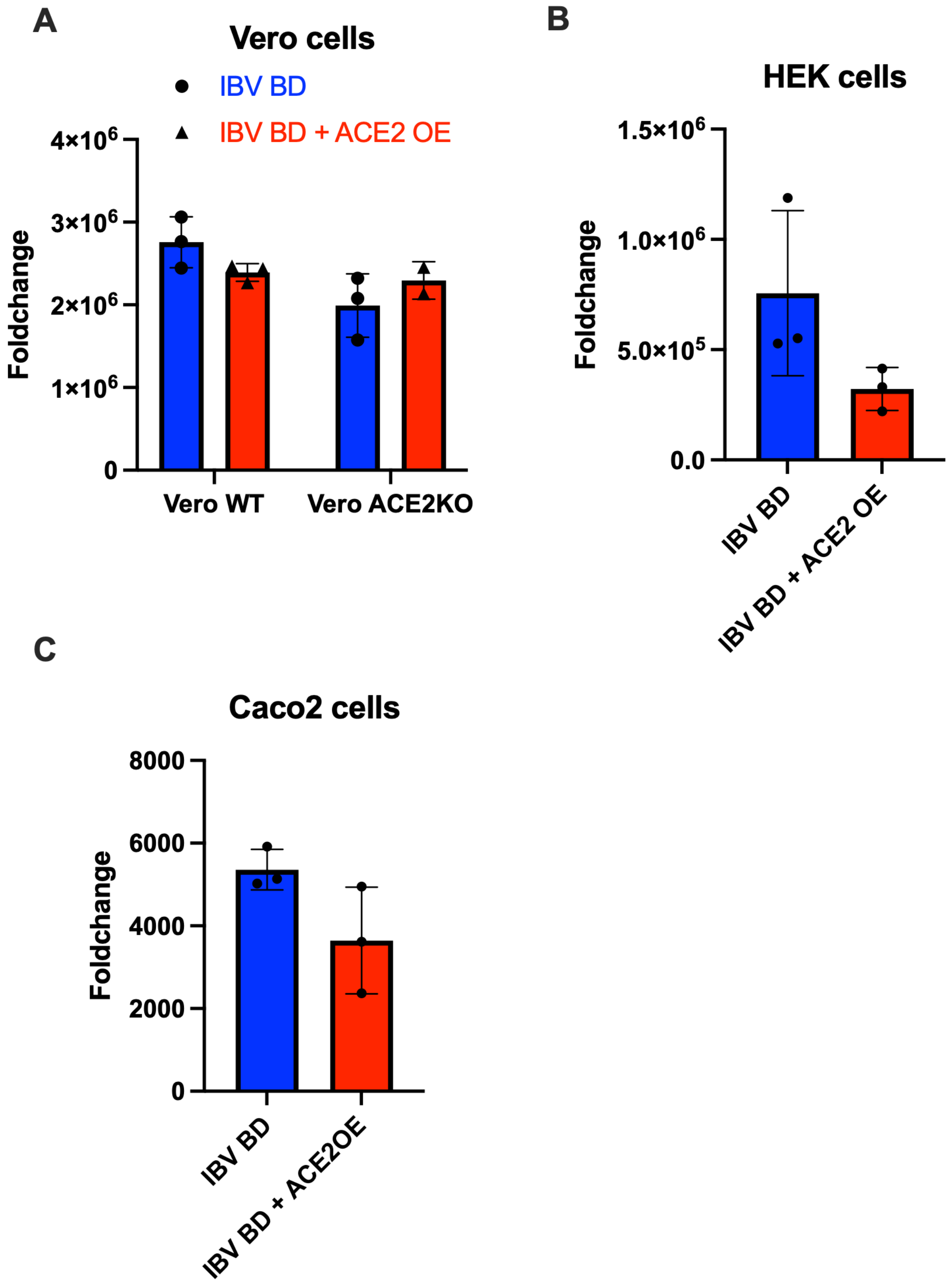


Supplementary Fig. S1 Evaluation of ACE2 overexpression during IBV infection in additional cell models. (A) WT and ACE2 KO Vero cells, (B) HEK293 cells and (C) Caco-2 cells, with or without ACE2 overexpression (ACE2 OE), were infected with IBV Beaudette (IBV BD). Viral RNA levels were quantified by RT-qPCR at 48 hours post-infection and are presented as fold changes relative to WT mock control. Bars represent the mean ± SD, and individual points represent biologically independent samples. Statistical significance was assessed using t tests, no statistically significant differences were detected.
